# New insights into the evolution of spider silk proteins illuminated by long-read transcriptomes

**DOI:** 10.64898/2026.02.03.703552

**Authors:** Kesen Zhu, Shiyi Zhou, Mo Lyu, Jiahao Xiang, Shaohan Niu, Yongping Huang, Lei Gao, Anjiang Tan, Hui Xiang

## Abstract

Spider silk has long fascinated scientists because of its exceptional mechanical properties, yet the evolutionary origins and diversification of spider silk proteins (spidroins) remain incompletely understood. By analyzing long-read spidroin transcripts from spiders representing key evolutionary lineages, we identify two ancestral proteins present in basal spiders: an alanine–serine-rich (AS-type) protein and a glycine–serine-rich (GS-type) protein. These ancestral proteins likely served as primary evolutionary templates for the diversification of modern spider silks. We show that the AS-type ancestral spidroin remained relatively conserved and ultimately gave rise to the specialized tubuliform spidroin (TuSp) used in eggcase silk. In contrast, the GS-type protein underwent extensive functional radiation, evolving into differentiated minor ampullate spidroins (MiSp) through the acquisition of distinct terminal domains and β-sheet–associated structural innovations. Our results further suggest that major ampullate spidroins (MaSp) may have originated from MiSp, with the GS motif representing an evolutionarily favorable genetic substrate for the emergence of high-performance silk proteins. Finally, we propose a revised evolutionary trajectory for flagelliform spidroins (Flag), suggesting that they were co-opted as components of ampullate silk during functional degeneration in modern non-web-weaving RTA clade spiders. Together, these findings provide a high-resolution framework for understanding the genetic innovations that drove the diversification of spider silk.

## Introduction

Beyond Lepidopteran silk, the dragline silk of spiders (order Araneae) has attracted considerable attention because of its exceptional tensile strength and toughness (Blamires et al., 2017). Unlike Lepidopterans, which typically produce a single silk type, spiders use specialized silk glands to spin a wide diversity of fibers. At the genetic level, this functional diversity reflects the evolutionary radiation of spider silk proteins (spidroins), which are encoded by a multigene family. In web-weaving spiders, dragline silk consists primarily of major ampullate (MA) silk, which is composed of major ampullate spidroins (MaSp).

In a typical orb web, MA silk forms the frame and radial threads (Dugger et al., 2020), whereas its counterpart, minor ampullate (MI) silk (Papadopoulos et al., 2009), provides additional structural reinforcement (Chen et al., 2012). MaSp and MiSp (particularly MaSp1 and MiSp) exhibit striking similarities in amino acid composition and repetitive motif organization (Nakamura et al., 2023; Papadopoulos et al., 2009). They are distinguished from other spidroins by their relatively short and variable repetitive units, in contrast to the long and highly stereotyped repeats characteristic of other silk types (Hayashi et al., 2004; Zhou et al., 2021). These include flagelliform spidroin (Flag) and aggregate spidroin (AgSp), which together form the “wet” spiral capture thread of araneoid webs (Stellwagen & Renberg, 2019), as well as Flag and cribellar spidroins (CrSp) in the “dry” capture threads of deinopoid webs (J. E. Garb et al., 2006). Additional specialized spidroins include the eggcase-associated tubuliform (TuSp) and aciniform (AcSp) spidroins (Garb & Hayashi, 2005), and pyriform spidroin (PySp), which is used to produce attachment discs (Chaw et al., 2017). Chemically, ampullate spidroins are enriched in alanine (A) and glycine (G), forming poly-A, poly-GA, or GGX motifs. In contrast, TuSp, AcSp, and PySp contain lower proportions of glycine but are enriched in serine (S) within their stereotyped repeats (Hayashi et al., 2004). Ampullate spidroins are further distinguished from Flag by C-terminal domain substitutions that facilitate the liquid-to-solid structural transitions required for fiber assembly (Rat et al., 2023).

The divergence of these specialized spidroins raises fundamental questions about how ancestral genetic material in basal spider lineages evolved into the diverse spidroins observed in modern spiders (Arakawa et al., 2022; Garb et al., 2010). Phylogenetically, Araneae comprise the basal Mesothelae, the subsequently diverged Mygalomorphae, and the highly diverse Araneomorphae (Craig, 2004). Araneomorphae include the orb-weaving Araneoidea and Deinopoidea, as well as the “RTA” (retrolateral tibial apophysis) clade, which represents the most successful non-web-building lineage derived from orb-weaving ancestors (Arakawa et al., 2022). Although MaSp is thought to have emerged during the initial radiation of Araneomorphae (Craig, 2004), the evolutionary origin of GA-rich ampullate spidroins remains unresolved (Fan et al., 2023). Efforts to identify ancestral forms in basal lineages have been limited by incomplete genomic resources. Although GA- and poly-A–rich motifs occur in Mygalomorphae, their relatively low glycine content suggests that they are unlikely to represent direct precursors of modern ampullate spidroins (Garb et al., 2007). In the basal Mesothelae, reported spidroin sequences are largely restricted to CySp-like and AcSp-like proteins (Arakawa et al., 2022; Starrett et al., 2012), further obscuring the nature of ancestral spidroins. In addition, the apparent loss of Flag spidroins in the RTA clade, which produces diverse silks for draglines and egg protection but lacks foraging webs (Arakawa et al., 2022), poses an intriguing evolutionary question regarding how silk components are lost or repurposed.

In this study, we utilized long-read transcriptomics to analyze 12 spider species representing both the most ancient and the most derived lineages of Araneae. We identify two primordial spidroins in basal Mesothelae that likely served as ancestral templates for modern AS-rich and GA-rich spidroins. Our analyses reveal contrasting evolutionary trajectories for these proteins, including strong conservation of the eggcase spidroin TuSp and extensive innovation of MiSp. We further propose that MiSp-like proteins provided the genetic foundation for the emergence of MaSp and identify poly-GS motifs as a versatile substrate for silk protein evolution. Finally, we present evidence that Flag spidroins were not simply lost in RTA spiders but were co-opted as auxiliary components of ampullate silk during their functional degeneration.

## Results

### Identification of spidroin resources across representative nodes in spider evolution

Using long-read transcriptome sequencing corrected with short-read RNA-seq data, we identified a total of 60 spidroin sequences across 12 representative spider (Araneae) species, among which 30 were full-length (figure supplements 3 and 4, Dataset 1). These species span the major evolutionary lineages of spiders, including Mesothelae, Mygalomorphae, Haplogynae, and the later-diverging Araneoidea and RTA groups within Araneomorphae. We constructed a phylogenomic tree based on single-copy orthologous proteins to resolve lineage relationships and estimate divergence times (Figure 1). The resulting topology and divergence estimates are generally consistent with the established phylogenetic backbone of spiders (Fernandez et al., 2018; Garrison et al., 2015). Notably, species from the families Eresidae and Araneoidea clustered together into a single clade that is sister to the RTA clade (Figure 1—figure supplements 1 and 2). We then mapped the identified spidroins onto the species tree (Figure 1). Possibly due to the limitation of whole-body sampling, we were not able to obtain the complete repertoire of spidroins for each spider. However, we were able to reconstruct a consistent evolutionary landscape, showing that basal spiders have fewer and more divergent spidroins, while the emergence of Araneomorphae is accompanied by diversification of spidroins.

**Figure 1.**
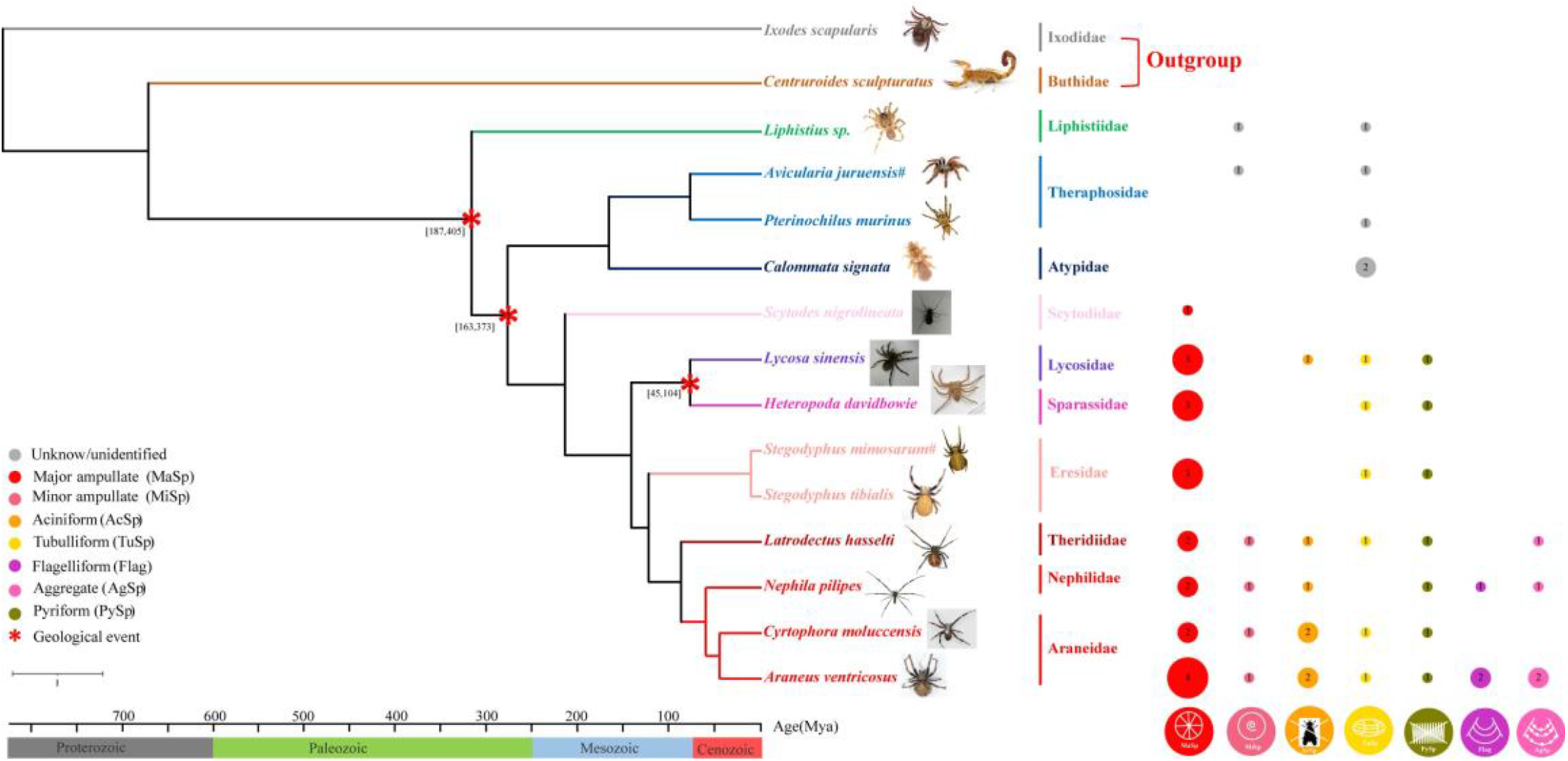
Phylogene and identified spidrioins of the spiders in this study. The phylogenetic status of the spiders was marked with different colors. the green line represents Mesothelae, the blue line represents Mygalomorphae, and the red and purple line represents Araneomorphae, specific species are marked at the end of the branch. An asterisk (*) indicates that an event with a significant impact is likely to occur, the ruler represents when the event occurred, the specific time scale is shown on the X axis in millions of years. Each colored box represents a silk type, the gray boxes represent unidentified spidroins, and numbers in each box indicate the total number of occurrences of each silk type.

### Two highly divergent putative spidroins in a basal species belonging to Mesothelae

In the basal Mesothelae genus *Liphistius*, which represents the earliest-diverging spider lineage (Wheeler et al., 2016; Xin et al., 2015), we identified two highly divergent spidroin-like transcripts encoding putative spidroin-like proteins, *Liphistius* sp._6705 and *Liphistius* sp._5400 (Figure 2A). As to *Liphistius sp*._6705, we recovered 12 repetitive units along with the C-terminal sequence. Each repetitive unit contains poly-alanine (polyA) and poly-threonine (polyT) motifs, while the remaining non-polymeric regions are dominated by alanine (A) and serine (S). We termed this the AS-type spidroin. The second transcript, *Liphistius sp*._5400, which we term the GS-type spidroin, was recovered as a full-length sequence and is characterized by a high frequency of glycine–serine (GS) repetitive motifs. The Poly-GS motifs are known β-sheet–forming elements and are also commonly observed in silk-producing insects, including silkworms (Mita & James, 1994), earwigs (Sutherland et al., 2010) and bagworm moths (Kono et al., 2019). In this protein, poly-GS motifs are interspersed with a small number of glycine–alanine (GA) sequences and terminate in a spacer region of unknown function, together forming a repetitive unit. We identified four such repetitive units together with the C-terminal sequence in this partial protein (Figure 2A). Notably, neither the N-terminal nor the C-terminal sequence of *Liphistius sp*._5400 shows homology to known spidroins (Figure 2B), suggesting that this spidroin-like protein may not assemble into typical silk fibers (Li et al., 2022), despite being substantially expressed (figure supplement 3).

**Figure 2.**
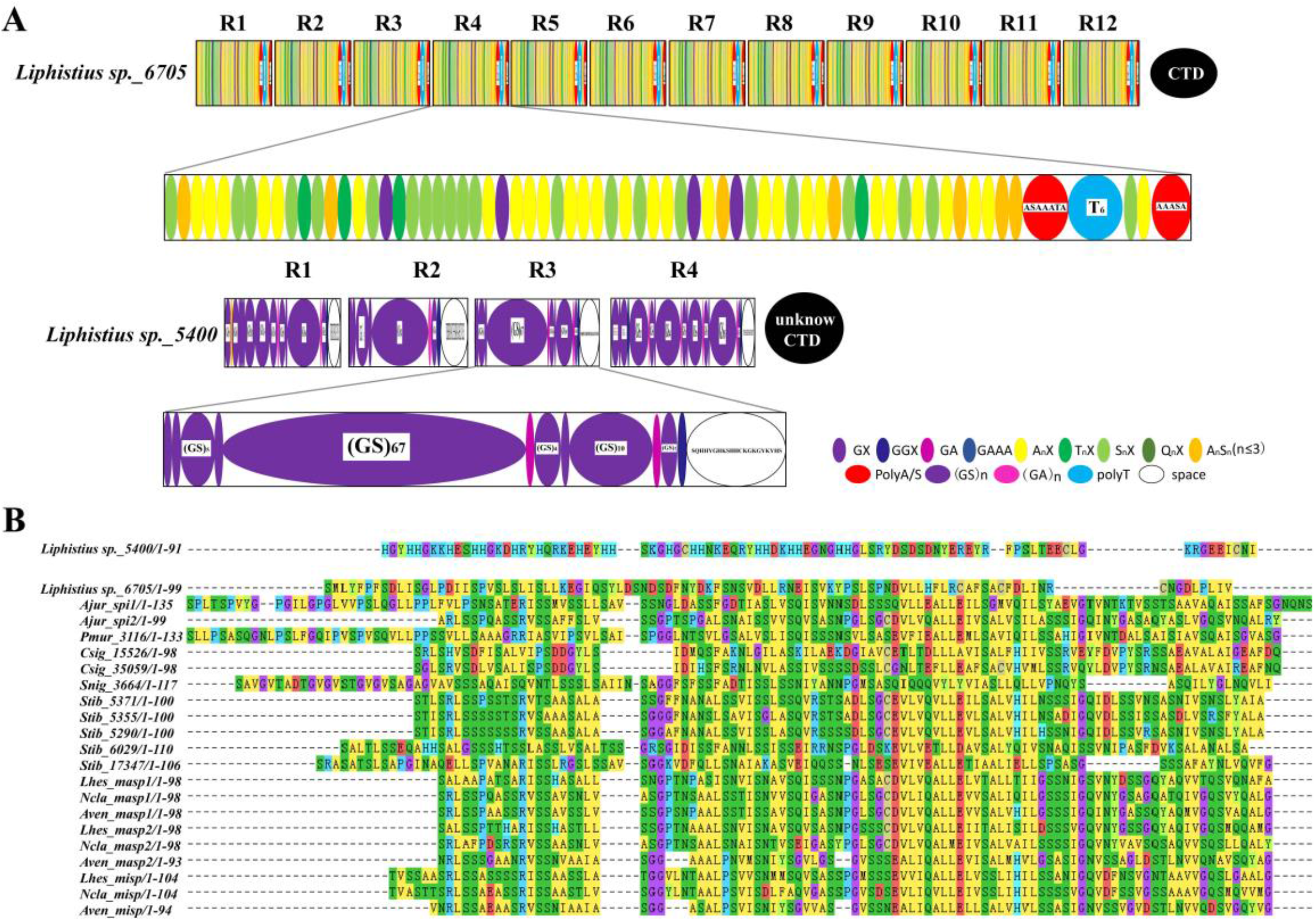
Analysis of silk protein structure of *Liphistius sp*. **(A)** Structural model analysis of the silk protein of *Liphistius sp*. from Thailand. (**B)** Alignment of amino acids in the C-terminal domain of spider silk protein sequences in multiple spider species, red arrows indicate the location of two ancient spider silk proteins. Ajur, *A. juruensis*; Pmur, *P. murinus*; Csig, *C. signata*; Snig, *S. nigrolineata*; Stib, *S. tibialis*; Lhes, *L. hesperus*; Ncla, *N. clavipes*; Aven, *A. ventricosus*.

### Phylogenetic trees based on different domains elucidated the evolutionary pattern of spidroins

Phylogenetic analyses based on different spidroin domains revealed distinct evolutionary patterns. In the N-terminal phylogeny, spidroins from the basal Mesothelae genus *Liphistius* (Xin et al., 2015) occupied the most basal position with robust support (Figure 3A). AgSp and PySp then diverged as a sister group to the remaining spidroins, suggesting that their terminal domains retain relatively basal characteristics. N-terminal sequences from the later-diverging Mesothelae genera *Heptathela* and *Ryuthela* (Xin et al., 2015) clustered with cribellar spidroin (CrSp), and subsequently with Flag (Figure 3A), indicating that spidroin differentiation may have occurred early in the earliest diverged spider lineage, Mesothelae. N-terminals from two Mygalomorphae species clustered with the two eggcase spidroins (AcSp and TuSp) and were parallel to the ampullate spidroin clade (Figure 3A). The N-terminal of a basal Haplogynae species within Araneomorphae is positioned as a sister lineage to the ampullate spidroin clade (Figure 3A). In the C-terminal phylogeny, *Liphistius* spidroins cluster with AgSp to form the basal clade (Figure 3B), further supporting the basal status of AgSp. In contrast, spidroins from Mygalomorphae are distributed across multiple clades, including TuSp, Flag, and the ampullate spidroins (Figure 3B).

**Figure 3.**
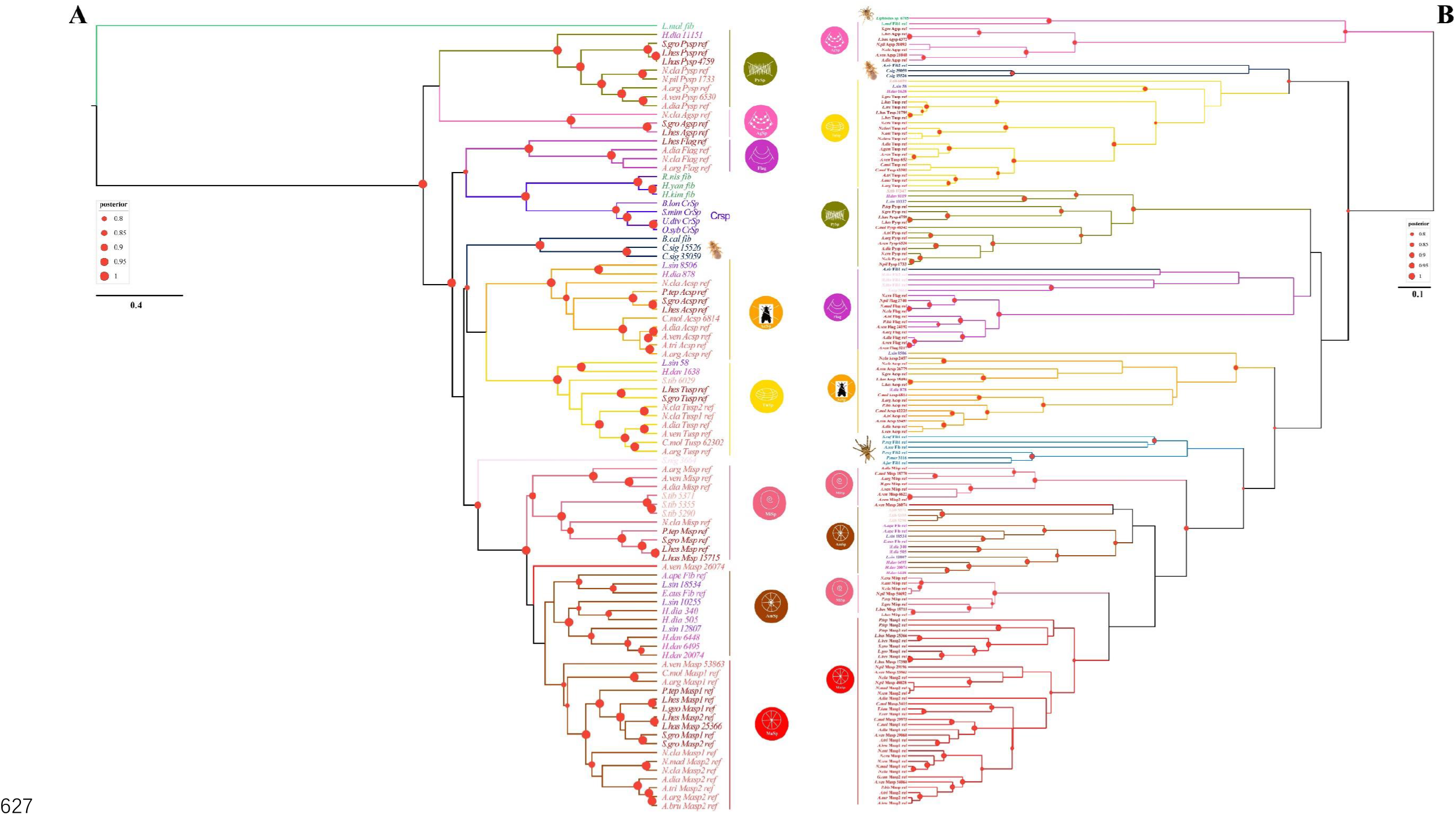
Bayesian Inference tree of spidroin amino-terminal regions. The colored icons represent different spidroin type, and the specific colour is used for each type. Colored icons surround spidroins categorized as flagelliform spidroin (Flag); aggregate spidroin (AgSp); pyriform spidroin (PySp); tubuliform spidroin (TuSp); aciniform spidroin (AcSp); major ampullate spidroin (MaSp); or minor ampullate spidroin (MiSp). At the end of the gene tree are various spider spidroin, which are represented by different colors and can be roughly divided into three categories: the green line represents Mesothelae, the blue line represents Mygalomorphae, the red and purple line represents Araneomorphae. Red dots indicate nodes with bootstrap values > 80%, Scale bar is substitutions per site. The primitive group Mesothelae and Mygalomorphae sequenced in this study are marked with photos. **(A)** Bayesian tree of spidroins N-terminal regions. **(B)** Bayesian tree of spidroins C-terminal regions.

Together, these phylogenetic trees indicate that spidroin terminal domains from *Liphistius* represent the most ancestral forms, with subsequent differentiation accompanying Mesothelae diversification. The emergence of Mygalomorphae coincides with substantial expansion and diversification of spidroins, while ampullate spidroins exhibit the greatest divergence in terminal domains. In contrast, phylogenetic analysis of repetitive units reveals less diversification in Mesothelae and Mygalomorphae compared with terminal regions (Figure 3), suggesting that innovation in repetitive sequences lagged behind the evolution of silk gland types. The repetitive unit phylogeny identifies a basal clade containing AS-type spidroins from Mesothelae together with multiple spidroins from Mygalomorphae (Figure 3). PySp and subsequently TuSp diverge from this group, forming basal clades relative to other spidroins (Figure 3). In contrast, the GS-type spidroin from *Liphistius* clusters with a spidroin from a Mygalomorphae species within the derived MiSp clade (Figure 4), implying that MiSp-like motif represents the basal form of ampullate spidroins. The repetitive unit tree further shows that MiSp clusters tightly with MaSp1, suggesting that MaSp1 may represent the ancestral copy within the MaSp family (Figure 4). Analyses of amino acid composition within repetitive motifs further clarify the divergence of the two major spidroin groups derived from the two types of Mesothelae spidroins, i.e., the AS-rich and GA-rich groups, respectively (Figure 4). The AS-rich group includes undifferentiated basal spidroins as well as the derived TuSp and PySp, whereas the GA-rich group comprises ampullate spidroins, particularly MiSp and MaSp1, together with the GS-type spidroin identified in *Liphistius*.

**Figure 4.**
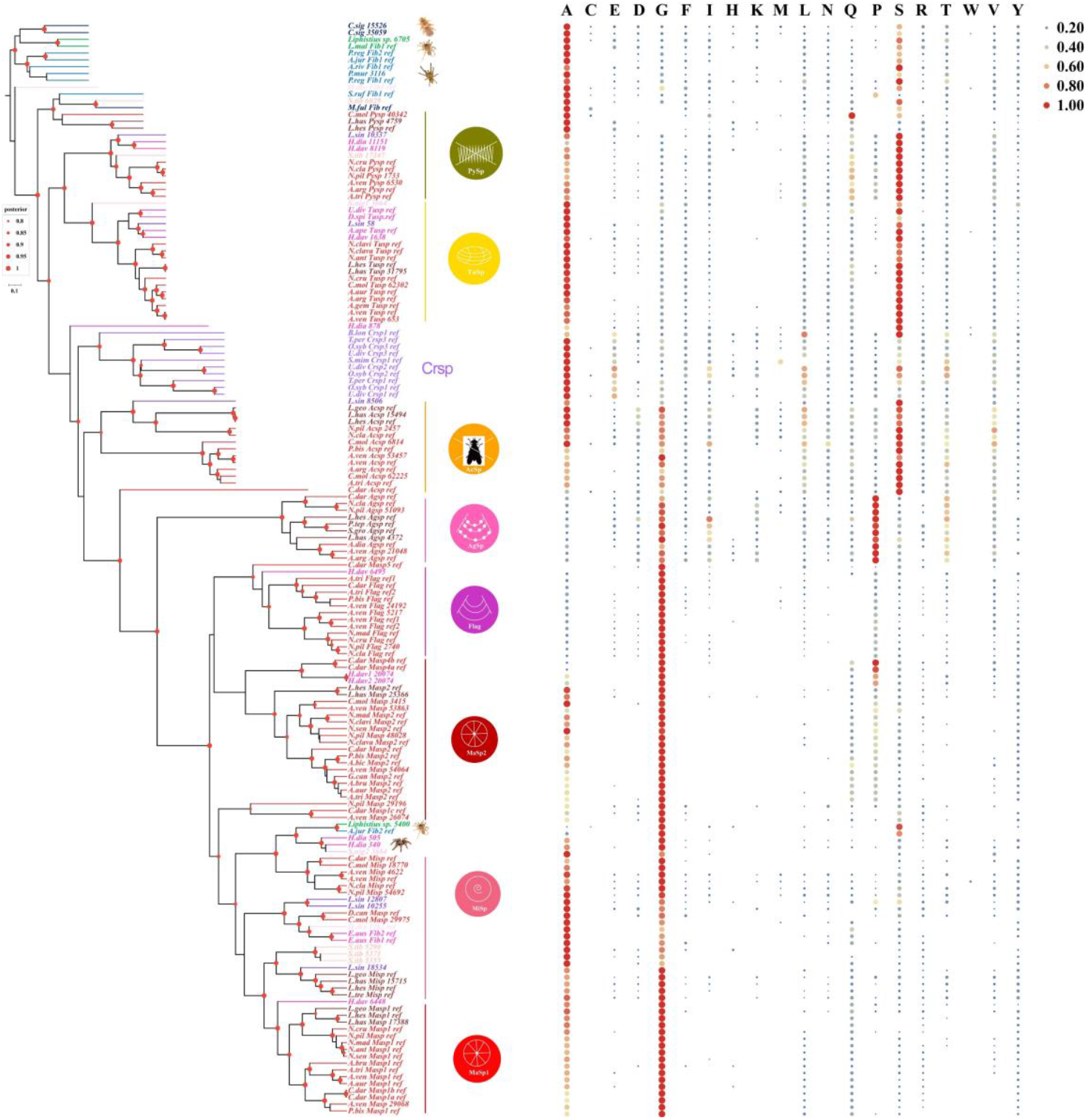
Bayesian Inference tree of spidroin repeat unit regions. The colored icons represent different spidroin type, and the specific colour is used for each type. Colored icons surround spidroins categorized as flagelliform spidroin (Flag); aggregate spidroin (AgSp); pyriform spidroin (PySp); tubuliform spidroin (TuSp); aciniform spidroin (AcSp); major ampullate spidroin (MaSp); or minor ampullate spidroin (MiSp). At the end of the gene tree are various spider spidroin, which are represented by different colors and can be roughly divided into three categories: the green line represents Mesothelae, the blue line represents Mygalomorphae, the red and purple line represents Araneomorphae. Red dots indicate nodes with bootstrap values > 80%, Scale bar is substitutions per site. The primitive group Mesothelae and Mygalomorphae sequenced in this study are marked with photos. Calculating the amino acid frequency from the repeating domain in spidroins and displaying it in a heatmap, the size of the circle and the depth of the color represent the frequency.

### Fairly conservation between the AS-rich ancestral spidroins and the egg-case TuSp

The repetitive units of AS-rich spidroins in Mesothelae and Mygalomorphae spiders were highly conserved, characterized by polyT and polyA motifs interspersed with other amino acids, as well as similar composition of additional motifs, such as AnX, SnX, and AnSn (where “X” represents any amino acid and “n” represents the number 1, 2, or 3) (Figure 5A). These features show substantial similarity with TuSp (Figure 5B), as all TuSp sequences also preserve polyT motifs, and one TuSp in RTA spiders additionally retains a polyA motif. TuSp proteins also show strong similarity to ancestral AS-rich spidroins in the composition of other repetitive motifs (Figure 5B). In contrast, polyA and polyT motifs are partially lost in the PySp of RTA spiders, which also feature novel spacer motifs in later-diverging spiders (Figure 5B), indicating greater divergence of PySp relative to TuSp.

**Figure 5.**
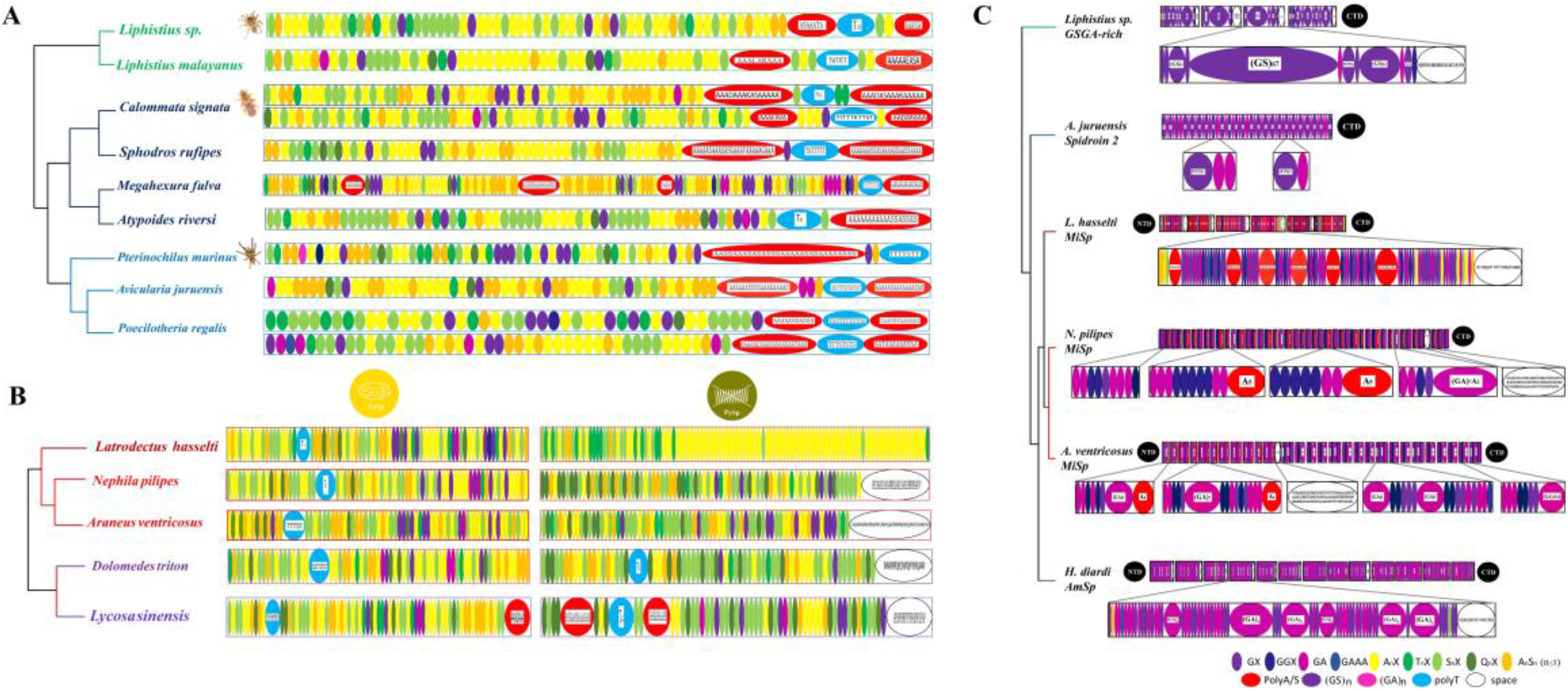
Evolutionary patterns of AS-rich and GSGA-rich ancient spidroin. Amino acid sequence pattern structure of spider silk is depicted, with different colored circles representing different mechanical properties of motif, the specific type motif has been annotated in the lower left corner of the figure. **(A)** The AS-rich spider silk sequence structure of Mesothelae and Mygalomorphae, with photos marking the primitive groups Mesothelae and Mygalomorphae sequenced in this study. **(B)** Tubuliform silk protein and pyriform silk protein patterns in Araneomorphae. **(C)** The pattern structure of GSGA-rich spidroin.

### The flagelliform-like spidroin in a basal RTA spider possesses MaSp-like terminal domains

Flagelliform spidroin (Flag) is a key component of prey-capture silk in typical orb webs and is commonly found in araneoid and cribellate orb-weaving spiders (Correa-Garhwal et al., 2022). Notably, we identified one Flag-like spidrion (*H*.*dav_6495*) in a basal RTA spider *Heteropoda davidbowie* (Wheeler et al., 2016) with highly homology to the Flag of two araneoid orb-web spiders, exhibiting substantial “GPGGX” and “GPG” motifs, which are typical features of Flag (Figure 6A). Transcriptome data confirmed its expression (figure supplements 5). This finding extends the existence of Flag to the non-web-weaving RTA clade. *Heteropoda* spiders are relatively close to cribellate orb-web spiders compared to other later-derived RTA spiders (Wheeler et al., 2016). Consistent with this, we did not identify Flag transcripts in the more derived RTA jumping spider *Hyllus diardi* (figure supplements 6). Genomic synteny analyses further indicate that Flag is absent from the araneoid outgroup *Stegodyphus dumicola* (Eresidae) as well as from the derived RTA species *Dolomedes plantarius* (Figure 6B). These results suggest that Flag may have originated independently in araneoid and cribellate orb-weaving lineages, arguing against a single origin of orb webs (Garb et al., 2006), and that Flag was subsequently lost during the evolution of RTA spiders.

**Figure 6.**
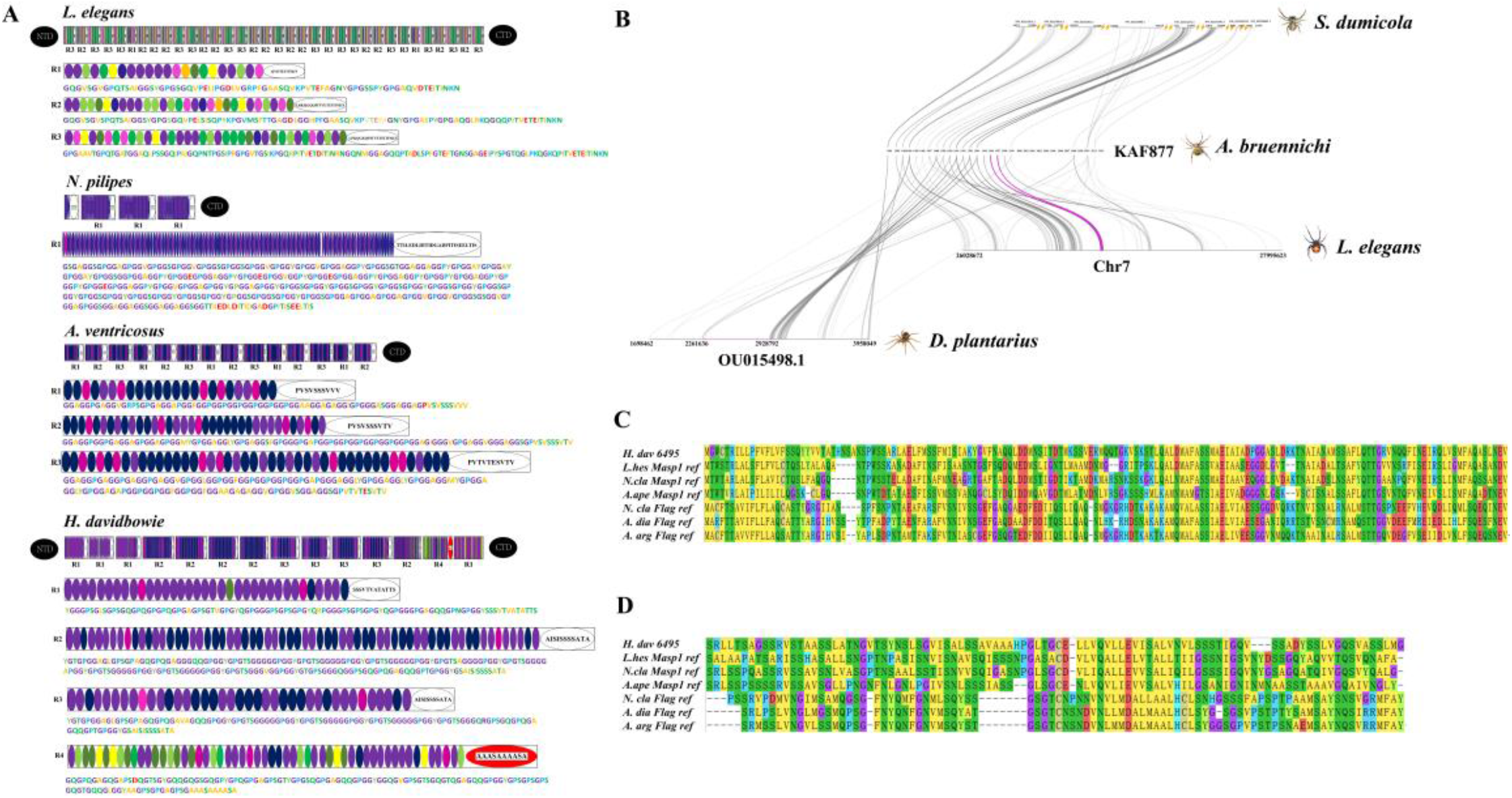
Analysis of important capture silk protein Flag sequence features. **(A)** Schematic diagrams of cob weaver (including repeat motif from *L. elegans*), two orb weavers (including repeat motif from *N. pilipes* and *A. ventricosus*) and RTA specialized clade spider (*H. davidbowie*). Each repeat is composed of different motifs which are shown as oval module labeled with different colors. Spacer, spacer region. NTD, N-terminal domain. CTD, C-terminal domain. **(B)** The collinearity of the upstream and downstream of *A. bruennichi* Flag gene in the ancient species of Araneomorphae (*S. dumicola*), the cobweb spider (*L. elegans*) and RTA specialized clade spider (*D. plantarius*) genome, the gray lines represent the genomic regions of other species compared to the Flag gene, the purple line represents the alignment position of the Flag gene. **(C)** N-terminal sequence alignment of MaSp1, Flag from 3 spiders of Araneoidea and *H. dav* spiders. **(D)** C-terminal sequence alignment of MaSp1, Flag from 3 spiders of Araneoidea and *H. dav* spiders.

Importantly, both the N- and C-terminals of this Flag-like spidrion (*H*.*dav_6495*) are highly homologous to those of ampullate spidroins rather than typical Flag (Figure 3 and Figure 6C, D). The N- and C-terminals function in maintaining high conservation of soluble spidroins in the corresponding silk glands and in the rapid assembly of spidroins (Andersson et al., 2014; Eisoldt et al., 2012). Specifically, the C-terminal of ampullate spidroins differs from that of Flag in that it undergoes structural transitions that contribute to greater toughness (Rat et al., 2023). Therefore, this switch of terminals from Flag-type to MaSp-type in the basal RTA spider implies that the Flag-like spidroin may have evolved to be adaptive to the microenvironment of the ampullate silk gland, adopting a MaSp-like assembly mechanism.

## Discussion

Supported by long-read transcript sequences combined with published data, our study provides a novel opportunity to decipher the evolutionary scenario of spider silk protein (spidroin) genes. Notably, in basal spider lineages, the N- and C-terminals of undifferentiated spidroins show greater diversification than the repetitive units (Figure 3 and 4). Both N- and C-terminals are important for silk storage and assembly, responding to the microenvironment of the silk gland (Andersson et al., 2014). Therefore, the phylogenetic relationships of spidroin terminal sequences may reflect the evolution of the corresponding silk gland microenvironments. The diverse phylogenetic positions of terminal sequences in these undifferentiated spidroins suggest that differentiation of silk gland microenvironments may have occurred earlier than genetic innovations in repetitive sequences. Supporting this idea, Mygalomorphae spiders possess morphologically complex silk-spinning apparatuses (Ferretti et al., 2017). Future in-depth molecular and genomic studies of silk glands in basal spiders will provide additional insight into these evolutionary dynamics.

In the basal-most lineage, i.e, *Liphistius sp* of Mesothelae (Wheeler et al., 2016; Xin et al., 2015), we discovered two genes with completely distinct characteristics: the AS-type spidroin and the GS-type spidroin-like gene. The coexistence of these two genes highlights the different evolutionary routes of dragline silk proteins (ampullate spidroins) and AS-rich spidroins (Fan et al., 2023). Notably, in the subsequently diverged basal lineage of Mygalomorphae, AS-type spidroins were well conserved, whereas the ancestral GS-type protein evolved into a typical spidroin. This divergent evolution may have occurred prior to the split between Mygalomorphae and Mesothelae.

The AS-type spidroins, rich in alanine and serine with mosaic poly-threonine and poly-alanine motifs, likely represent the earliest spidroins. This preliminary form of AS-rich spidroins was already used for silk production at the emergence of spiders. Sequence analysis suggests that the AS-rich TuSp is highly conserved with the AS-type spidroin in *Liphistius sp*, extending the proposed conservation of TuSp to the most basal spider lineages (Garb et al., 2007). Broad conservation of TuSp has also been reported in Tengella perfuga, a cribellate spider from the RTA clade (Correa-Garhwal et al., 2018), further supporting its evolutionary conservation. TuSp is a pivotal protein in spider silk evolution, constituting protective silk that forms the outer layer of the spider egg sac, which is crucial for offspring survival. Mesothelae and Mygalomorphae spiders also produce eggcase silk (Palmer et al., 2010). Strong selective constraints on this AS-rich spidroin may contribute to its functional stability.

The polyGS-bearing protein (GS-type) is suggested to be the prototype of GA-rich ampullate spidroins. It appears not to have been initially used for silk production, given its non-homologous N- and C-terminal sequences relative to canonical spidroins (Figure 2). In the subsequently diverged Mygalomorphae lineages, a novel spidroin evolved from this genetic material through adoption of canonical spidroin terminal sequences. We also demonstrate that the spidroin in the mygalomorph spider *Avicularia juruensis* is MiSp-like instead of MaSp2-like (Bittencourt et al., 2010). Fibers composed of these polyGS-bearing proteins are relatively weak in tensile strength (Collin et al., 2009). Along spider evolution, increases in GA motifs and further recruitment of polyA motifs in these silk proteins enhanced mechanical properties, as observed in the evolved MiSp of web-weaving and RTA spiders [3, 5]. Phylogenies of spidroin terminal sequences indicate a derived position of MiSp and MaSp, with MaSp appearing to have differentiated later than MiSp (Figure 3). Combining this with repetitive sequence analyses, we propose that MaSp (the dragline spidroin) originated from MiSp, potentially before the emergence of the basal Araneomorphae lampshade spider *Hypochilus thorelli*, which possesses MaSp (Starrett et al., 2012).

GS-rich and derived GA-rich silk proteins are also widely distributed in silk-producing insects. For instance, in the basal insect lineage Embioptera (webspinners), polyGS motifs are used in silk proteins (Collin et al., 2011). Later-diverged Lepidoptera caterpillars also spin MiSp- or MaSp-like silks in early larval stages (figure supplements 7), and their silk glands show similarities to spider ampullate silk glands (Hu et al., 2023; Kono et al., 2019). Although more evidence is needed to clarify potential convergent or parallel evolution between spider ampullate spidroins and Lepidopteran silk fibroins, GS-rich genetic material appears to be evolutionarily favorable for the formation of modern silk proteins.

Interestingly, we identified a novel full-length spidroin gene in the basal Araneomorphae species *Scytodes nigrolineata* (*S. nig_3664*) (figure supplements 8A). This highly expressed spidroin contains six repetitive units, each comprising two tightly connected motifs: one AS-rich and one GA-rich, representing the two types of spidroin composition (Figure 4—figure supplements 8). The AS-rich motif is highly similar to TuSp, whereas the GA-rich motif resembles MiSp (figure supplements 8). This finding highlights vigorous rearrangement of spidroins during this evolutionary period and underscores the complexity of silk protein evolution. Further genomic evidence will help clarify the nature of these rearrangements. The same confusion may be the case of the Flagelliform spidroin (Flag). Its origin could not be traced through sequence analysis, but its evolutionary scale was extended to the non-web-weaving RTA lineage. The origin of the RTA clade from the ancestor of Deinopoidea orb-web spiders (Arakawa et al., 2022; Yu et al., 2022) coincided with degradation of web-weaving capacity. As a typical web component, flagelliform silk functions as capture thread. Flag is a high-cost spidroin and thus poses a selective burden during adaptive evolution of RTA spiders. In contrast, ampullate silks and spidroins are universally essential as dragline and lifeline silk for spiders and even some caterpillars. By examining the Flag-like spidroin in basal RTA spiders, we observed that its terminals are possibly adaptive to the ampullate silk gland. Given that later RTA spiders lost Flag, we propose that it may have initially been incorporated into ampullate silks in early RTA spiders, followed by progressive degradation and eventual loss of the corresponding gene in derived RTA species.

## Materials and methods

### Spider samples

Through field investigations, we initially collected eight species of spiders based on morphological characteristics in four geographically distinct regions in China (Sichuan, Yunnan, Xinjiang, Guangdong). These species include: *Calommata signata Karsch* (Mianyang, Sichuan: 31.30°N, 104.42°E), *Scytodes nigrolineata* (Liangshan, Sichuan: 27.89°N, 102.26°E), Stegodyphus tibialis (Dali, Yunnan: 25.34°N, 100.13°E), *Latrodectus hasselti* (Liangshan, Sichuan: 27.89°N, 102.26°E), *Lycosa sinensis* (Yili, Xinjiang: 43.55°N, 81.20°E), *Hyllus diardi* (*Xishuangbanna*, Yunnan: 22.01°N, 100.80°E), *Nephila pilipes* (Luoping, Yunnan: 24.884626°N, 104.31°E), and *Cyrtophora moluccensis* (Liannan, Guangdong: 24.77°N, 112.28°E). Additionally, through donations from spider enthusiasts, we collected three species: *Liphistius sp*., *Pterinochilus murinus, Heteropoda davidbowie*. All spider samples were collected and photographed, as shown in Figure 1. Spiders used for high-throughput sequencing were starved for 2 days to prevent exogenous contamination and then stored at −80°C after rapid freezing in liquid nitrogen.

### RNA extraction, long read transcriptome sequencing (ISO-Seq) and Illumine RNA-seq

Spider samples were sent to Novogene Co, Ltd for RNA extraction and transcriptome sequencing. Briefly, the whole body of spider samples was ground in liquid nitrogen, and total RNA was extracted using TRIzol reagent (Invitrogen) on dry ice. RNA concentration was accurately quantified using Qubit (Invitrogen Life Technologies, USA), while the integrity of the RNA was detected using an Agilent 2100 Bioanalyzer (Agilent Technology, USA). For ISO-Seq, libraries were prepared using the PacBio Iso-Seq library preparation method, followed by sequencing on the Pacific Biosciences Sequel II platform at the University of Maryland Institute for Genome Sciences. For RNA-Seq, libraries were prepared using the Illumina RNA-seq library preparation method and sequenced on the Illumina HiSeq™ 2000 platform.

### Raw data processing

Raw data were filtered and processed using SMRTlink v6.0 software with the following parameters: --minLength 50, --maxLength 15000, --minPasses 2, --min_seq_len 200. Subreads sequences were obtained, and circular consensus sequences (CCS) were generated by correcting subreads sequences. CCS containing 5’-primer, 3’-primer, and polyA were detected. CCS were classified into full-length non-chimeric (FLNC) sequences and non-full-length (nFL) sequences. FLNC sequences of the same transcript were clustered using the ICE algorithm to obtain consensus sequences, which were then corrected by nFL sequences using Arrow software. Sequences were further corrected using Illumina RNA-seq reads with LoRDEC v0.7 (Salmela & Rivals, 2014) software, followed by de-redundancy processing using CD-HIT-EST v4.6.8 (Li & Godzik, 2006) (-c 0.95). The resulting non-redundant transcripts were used for subsequent analyses. Long-read transcriptome data (SRR12208061) and Illumina RNA-seq data (SRR11905576) of A. ventricosus were downloaded from the NCBI SRA database and analyzed using the same pipeline. The average length of sequences and contig N50 size were calculated (figure supplements 1 and 2).

### Annotation of genes

The genes were functionally annotated by homology-based searches against the following database: NCBI Non-redundant Protein (NR), NCBI Non-redundant Nucleotide (Nt), SwissProt, Kyoto Encyclopedia of Genes and Genomes (KEGG), euKaryotic Ortholog Groups (KOG) and Gene Ontology (GO).

### Construction of pseudo-coding genome

Non-redundant transcripts were assembled to identify full-length genes (CDSgenome) using Cogent v3.2 software. Each transcript family was folded into one or several unique transcript models (UniTransModels), and redundant isoforms were further collapsed using minimap2 (Li, 2018) alignment and cDNA_Cupcake v5.3. The constructed pseudogenome was analyzed with ANGEL v2.4 (Shimizu et al., 2006) software (--min_aa_length 100) for Open Reading Frame (ORF) prediction analysis, and BUSCO v5.0 (Simão et al., 2015) software was used to evaluate data integrity after processing.

### Orthologous gene identification

Genome information files of the spider *Stegodyphus mimosarum* (GCA_000611955.2) and two outgroup species Ixodes scapularis (GCA_016920785.2) and *Centruroides sculpturatus* (GCA_000671375.2) were developed. Single copy orthologous proteins were identified by reciprocal blast (Mount, 2007) (evalue cutoff: 10-5), with the gene set of *S. mimosarum* as a reference, against the annotated gene set of each of the 14 species. All single-copy orthologs of each species were aligned using Mafft v7.407 (Katoh et al., 2002) software and concatenated into one super gene for each species.

### Phylogeny reconstruction and divergence time estimation

The well-aligned single-copy orthologs were subjected to Maximum likelihood–based phylogenetic analysis using RAxML v8.2.12 (Stamatakis et al., 2005) with the following parameters: -f a -T 12 -p 12345 -x 12345 -m PROTGAMMAAUTO -# 1000. Divergence time was calculated using the MCMCtree program in the PAML v4.8 (Yang, 2007) package, and fossil records obtained from the TimeTree database (Kumar et al., 2022) were used for result calibration.

### Identification and preliminary analyses of spidroins

Released spidroin sequences in the NCBI NR database were collected, and representative complete spider silk protein sequences were used as reference sequences for blastp (Mount, 2007) (e-value cutoff: 10-5) against the protein set of each of the 12 spider species. Fitted sequences were manually checked for a typical spider silk protein characteristics, shorter sequences or missense mutations. Error-free long spider silk sequences were verified by blastp against the NR database. Identified spidroin sequences were analyzed for signal peptide prediction using SignalP v5.0 and hydrophobicity prediction using Protscale and Protparam tools in the ExPASy platform (https://www.expasy.org/). Selected error-free spidroin sequences were divided into three regions: N-terminal, repeat region, and C-terminal. Amino acid frequency of repetitive sequences was calculated and a heatmap was generated using Tbtools v2.056 (Chen et al., 2020) software.

### Phylogenetic analysis of spidroins

The lack of homology of the terminals Lera_5400 to any other spidrion supports the ancient status of Liphistius spidrions. We excluded these sequences and investigated the phylogenetic relationship of amino acid sequences of the rest spidroins N-terminal, C-terminal sequences, and repetitive motifs, respectively, with some publicly available sequences added in (figure supplements 5). The N-terminal, repeat region, and C-terminal of spidroins were aligned separately using MEGA v7 (Kumar et al., 2018). Aligned sequences of each region were utilized to construct gene trees using beast v1.10 (Suchard et al., 2018) software with the Bayesian method and the following parameter settings: chain length of 20,000,000, logEvery of 1000, and burnin of 0.1.

### Calculation and visualization of spider gene expression

The Minimap2 (Li, 2018) software was used to compare the redundant transcript data of the full-length transcriptome to the pseudo-coding genome constructed by Cogent v3.2 software, the bowtiev2 (Langmead & Salzberg, 2012) software was used to compare the raw data of second-generation transcriptome sequencing to the pseudo-coding genome. Use the Samtools v0.1.19 (Li et al., 2009) tool to convert the sam format generated after the comparison into a bam format file, the bam files are sorted and indexed, and then the bam files are imported into the IGV software for visual analysis.

## Data availability

All data needed to evaluate the conclusions in the paper are present in the paper and/or the Supplementary Materials. All raw data was deposited at the NCBI in the sequence read archive (SRA) under accession number (BioProject Number: PRJNA754522).

## CRediT authorship contribution statement

H. X. conceived and designed the investigation. K.S.Z., G.L., M.L and J.H.X. collected and prepared the samples. K.S. Z. analyzed the data. K.S. Z. and H. X. wrote the paper. A.J. T., H. X, and Y.P. H. revised the manuscript. All the authors read and approved the final manuscript.

## Acknowledgement

We thank Prof. Wen Wang for help in revised the manuscript and give suggestions for this work. We thank Prof. Sheng Li for insightful comments. We thank Ms. Jiajia Zhang for her help in handling the spiders.

